# Bacterial lipopolysaccharide modulates immune response in the colorectal tumor microenvironment

**DOI:** 10.1101/2022.04.26.489473

**Authors:** A.K. Sulit, M. Daigneault, E. Allen-Vercoe, O.K. Silander, B. Hock, J. McKenzie, J. Pearson, F.A. Frizelle, S. Schmeier, R. Purcell

## Abstract

Immune responses can have opposing effects in colorectal cancer (CRC), the balance of which may determine whether a cancer regresses, progresses, or potentially metastasizes. These effects are evident in CRC consensus molecular subtypes (CMS) where both CMS1 and CMS4 contain immune infiltrates yet have opposing prognoses. The microbiome has previously been associated with CRC and immune response in CRC but has largely been ignored in the CRC subtype discussion. Using CMS subtyping, we aimed to determine the contributions of the microbiome to the pleiotropic effects evident in immune-infiltrated subtypes. We integrated host gene-expression and meta-transcriptomic data to determine the link between immune characteristics and microbiome contributions in these subtypes and identified lipopolysaccharide (LPS) binding as a potential functional mechanism. We identified candidate bacteria with LPS properties that could affect immune response, focusing on *Fusobacterium periodonticum* and *Bacteroides fragilis* in CMS1, and *Porphyromonas asaccharolytica* in CMS4. Treatment of peripheral blood mononuclear cells (PBMCs) with LPS isolated from these bacteria showed that *F. periodonticum* stimulates cytokine production in PBMCs while both *B. fragilis* and *P. asaccharolytica* had an inhibitory effect. Furthermore, LPS from the latter two species can inhibit the immunogenic properties of *F. periodonticum* LPS when co-incubated with PBMCs. We propose that different microbes in the CRC tumor microenvironment can alter the local immune activity, with important implications for prognosis and treatment response.

## Introduction

Colorectal cancer (CRC) tumors consist of a complex microenvironment whose characteristics affect the tumor’s progression, prognostics, and therapy response (1,2). The immune-cell infiltrate within the microenvironment plays a key role in CRC as it can either enhance or inhibit tumor development. The induction of a tumor-directed immune response can result in the influx of effector cells and subsequent tumor cell death. However, in other contexts, the tumor itself may subvert the immune response and rather than eliminating the tumor, immune infiltrates may contribute to chronic inflammation and provide signals for cell growth and vascular changes (2,3). Different cytokines and T-cell subsets have been shown to be capable of both promoting or inhibiting cancer progression, and this likely reflects differences in the immunoregulatory signals provided by different tumor microenvironments.

Due to CRC heterogeneity, several different subtyping classifications have been proposed (4–9). Two subgroups are commonly identified among these different schemes, one characterized by microsatellite instability (MSI) and immune activation, and the other defined by angiogenic and mesenchymal characteristics (7). The Colorectal Cancer Subtyping Consortium combined data from several large subtyping studies to generate a classification system of four consensus molecular subtypes (CMS), based on gene expression data (5). Subsequent studies of the immune phenotypes associated with each of these molecular subtypes revealed substantive differences in the composition of the immune infiltrates, with the good prognosis CMS1 (MSI-Immune) being immunogenic and the poor prognosis CMS4 (mesenchymal) being inflamed and characterized by immunosuppression (10–12).

Understanding immune responses in CRC is not complete without consideration of the microbiome, as evidenced by lower tumor susceptibility in germ-free rats compared to conventional rats upon carcinogen introduction (13,14); differences in tumor susceptibility between mice with different microbiome communities (15); and differences in microbiomes of cancers with deficient or proficient mismatch repair functions (16). Bacteria such as *Fusobacterium nucleatum, Porphyromonas asaccharolytica*, and *Parvimonas micra* have been identified as potential biomarkers in studies of CRC (17,18). Taxa associated with the oral cavity have also been found to be enriched in CRC (18–21), possibly driving the increase seen in species richness (18). Of the microbial functional pathways that have been associated with CRC, many involve immune responses. *F. nucleatum* subspecies *animalis* can induce CCL20, a chemokine that plays a role in recruitment of Th17, regulatory T-cells, and dendritic cells (22), while Enterotoxigenic *B. fragilis* (ETBF) toxin is associated with murine colon tumor formation, through activation of signal transducer and activator of transcription-3 (STAT3) with Th17 responses, also involving IL-17 and IL-23 (23).

The CMS1 and CMS4 subtypes differ with respect to both their immune composition and prognoses. Although the microbiome is known to be a strong modulator of immune responses, its potential role in driving the differing immune compositions of CMS1 and CMS4 subtypes is yet unexplored. In this study we analyzed the microbiome composition in each subtype and detected differences in microbiome signatures associated with different patterns of immune activation in CMS1 and CMS4. We therefore aimed to determine how microbes from different subtypes of CRC tumors could potentially affect the immune environments characteristic of these respective tumors.

## Methods

### Sample Collection and Handling

A total of 308 samples were collected during surgical resection of colorectal tumors from patients who had not received chemotherapy prior to surgery. Patients with diagnosed HNPCC or FAP were excluded. All participants provided written, and informed consent and the study was approved by the University of Otago Human Ethics Committee with approval number ***H16/037***. During surgery, samples were taken, frozen in liquid nitrogen, and stored at −80°C. Before RNA extraction, samples were first equilibrated for at least 48 hours at −20°C in RNAlater ICE^™^ (Qiagen).

RNEasy Plus Mini Kit (Qiagen) was used to extract RNA from 15-20mg of tissue, disrupted using a Retsch Mixer Mill, including a DNAse treatment step in the procedure. Purified RNA was quantified using a NanoDrop 2000c spectrophotometer (Thermo Scientific, Asheville, NC, USA) and subsequently stored in −80°C.

### RNA Sequencing

Library preparation for RNA sequencing was carried out using the Illumina TruSeq Stranded Total RNA Library preparation kit (Illumina), with ribosomal RNA depletion using Ribo-Zero Gold. The Illumina Hi-Seq 2500 V4 platform was used for RNA sequencing, producing 125 bp paired-end reads. Each sample library was split into lanes to avoid technical bias and were later merged during the data processing phase. Merged raw data can be found under Bioproject ID PRJNA788974 in the NCBI SRA database.

### Consensus Molecular Subtype Classification

Fastq-mcf from ea-utils (24,25) and SolexaQA++ (26) were used for quality control and trimming of reads before merging of sequences from different lanes. Salmon (27) was then used to quantify transcript expression. The publicly available CRC CMS classifier (5) was used to categorize samples into one of four CMSs. Of the 308 samples, 260 were classified into a CMS, with 60 samples in CMS1, 145 in CMS2, 38 in CMS3, and 17 in CMS4. For subsequent analysis, we focused on the 260 classified samples.

### Bioinformatics Analysis

After trimming, the 260 samples were run through the MetaFunc pipeline (28) using default settings, except for setting reverse stranded option for featureCounts (29), and a species needing at least 0.01% abundance in at least one of the 260 samples to be included in the microbiome analysis. Databases used were those provided in https://metafunc.readthedocs.io/en/latest/usage.html#databases.

### Differential Expression and Gene Set Enrichment Analysis in Host

From the results of the MetaFunc analysis, we gathered the host-gene expression raw counts table into a matrix for input into *DESeq2* (30) with metadata information on their respective CMS. Using the 260 samples that had been classified into a subtype, *DESeq2* was used to obtain differentially expressed genes (DEGs) in one CMS compared to the average of the other three subtypes. Genes were considered differentially expressed if their Benjamini-Hochberg (BH) adjusted p-values were <0.05. Raw p-values and log-fold change generated through *DESeq2* were then used for ranking and sign information, respectively, in gene set enrichment analysis (GSEA) using the *clusterProfiler* package (31) with the C5 Ontology Gene Sets collection (version 7) from the molecular signatures database (MSigDB) (32,33). We considered a gene set enriched in a subtype if it had adjusted p-values of <0.05 and positive Normalized Enrichment Score (NES). We interrogated the enriched gene sets in the CMS subtypes using immune-related keywords as follows: “IMMUN”, “T CELL”, “INTERFERON”, “CYTOKINE”, “TOLL LIKE”, “LYMPHOCYTE”, “LEUKOCYTE”, “PATTERN RECOGNITION”, “LIPOPOLYSACCHARIDE”, “MHC”, “INFLAMMATORY”, “ANTIGEN”, “INTERLEUKIN”, and “BACTERIA”.

Detailed host analysis may be found at https://gitlab.com/alsulit08/2021_uoc_massey_lps-crc/-/tree/master/Bioinformatics.

### Differential Abundance of Microbes in the Microbiomes of CRC Subtypes

From the results of the MetaFunc analysis, raw counts of microbial taxonomies were gathered into a Phyloseq object (34), with metadata information on their respective CMS. *DESeq2* was used to identify differentially abundant microbes in either CMS1 or CMS4 compared to the average of the other three subtypes. Microbes were considered differentially abundant in a CMS if they had a log2 fold change > 0 and adjusted p-value < 0.05.

### Lipopolysaccharide-associated Bacteria

MetaFunc produces a table that indicates which bacterial taxonomy IDs have proteins that are annotated with gene ontology terms. All bacterial species with proteins annotated with “lipopolysaccharide biosynthetic process” or “lipid A biosynthetic process” were obtained for CMS1 and CMS4 samples, and then cross-referenced with differentially abundant microbes in CMS1 or CMS4, respectively, to obtain a list of differentially abundant bacteria that have proteins annotated with LPS-related processes.

### Lipopolysaccharide from Bacterial Strains

LPS was extracted from strains of *Fusobacterium periodonticum* (*1/1/54 (D10), 2/1/31*, and *1/1/41 FAA*), *Bacteroides fragilis* (*3/2/5, 2/1/16*, and *2/1/56 FAA*), and *Porphyromonas asaccharolytica* (*CC44 001F, and CC1/6* F2) using Bacterial Lipopolysaccharides (LPS) Extraction Kit (Alpha Diagnostic International, Catalog # 1000-100-LPS) as per the manufacturer’s instructions, resulting in a final yield of 30 μg/mL of LPS.

### PBMC Treatment with LPS from Bacterial Species

PBMCs (2×10^5^ cells) were incubated with LPS preparations (at least 16h) of varying concentrations (600 ng/mL, 60 ng/mL or 6 ng/mL) from *B. fragilis* (*3/2/5, 2/1/16*, and *2/1/56 FAA*), *F. periodonticum* (*1/1/54 (D10), 2/1/31*, and *1/1/41 FAA*) or *P. asaccharolytica* (*CC44 001F, and CC1/6* F2) at concentrations of 600, 60, or 6 ng/mL. For co-incubation tests, we used *B. fragilis* strain *2/1/16, F. periodonticum* strain *2/1/31*, and *P. asaccharolytica* strain *CC1/6 F2*. We treated the PBMCs with 6 ng/mL of *F. periodonticum* (strain *2/1/31)* LPS and 600 ng/mL of *B. fragilis* (strain *2/1/*16) or *P. asaccharolytica* (strain *CC1/6 F2*) LPS. As no-treatment controls, PBMC medium (RPMI, 10%FCS, 1% glutamine, 0.2% Penicillin/Streptomycin) or RPMI alone was added to the initial culture of PBMCs. Co-incubation experiments were conducted at least three times. For each repeated experiment, PBMCs were obtained from a different individual.

### Measurement of Cytokine Production and Statistical Analysis

Secreted cytokine expression was measured using LegendPlex Human Inflammation Panel 1 (Cat no. 740809) on a Beckman Coulter Cytomics FC500 Flow Cytometry Analyzer, following the manufacturer’s instructions. For all runs, baseline values of the cytokines from untreated PBMCs were obtained. Legendplex Software (Windows version 8 or MacOS version 7.1, using the 5-parameter curve fitting model) was used to assess final concentrations of the cytokines of interest. Paired student’s t-tests were used to test for differences in cytokine production between baseline PBMC and *F. periodonticum* treatment, and *F. periodonticum* treatment and *F. periodonticum* with either *P. asaccharolytica* or *B. fragilis* treatment. Effect Sizes were obtained using Cohen’s d, with Hedges correction, to account for small sample sizes.

Detailed analyses may be accessed at https://gitlab.com/alsulit08/2021_uoc_massey_lps-crc/-/tree/master/LPS_Experiments

## Results

### CMS1 and CMS4 have Enriched Gene Sets Involved in Immune Response

In order to compare which gene sets were enriched in different subtypes of CRC, we first used DESeq2 to compare gene expression in CMS1 samples with the average of the CMS2, CMS3 and CMS4 samples. There were 4736 genes that were significantly over expressed (adjusted p-value < 0.05) in CMS1 compared to the other subtypes. We carried out gene-set enrichment analysis (GSEA) on the list of genes as described in the methods section, and obtained those with positive normalized enrichment scores (NES) as gene sets enriched in CMS1. We obtained a total of 318 gene sets that had positive NES in CMS1, with adjusted p-value < 0.05. Several of the most enriched gene sets were related to immune responses (cytokines, antigen processing and presentation, and cell killing processes) as well as nuclear organization and replication processes. We therefore subset the enriched gene sets in CMS1 using immune-related keywords (see **Methods**). As it has been theorized that microbes might affect the balance of immune responses in the tumor microenvironment (TME), we sought to identify if response to microbes is captured among the enriched gene sets of our subtypes by including “BACTERIA” in the keywords.

We performed the same analyses for CMS2, CMS3, and CMS4, against the average of the other three subsets. We obtained no enriched gene sets using the immune-related keywords above in CMS2 and CMS3, consistent with previous studies describing these as “immune-neglected” (35). There were 2481 DEGs in CMS4 compared to the other subtypes. The top enriched gene sets for CMS4 primarily comprised terms corroborating its epithelial-mesenchymal-transition (EMT) and angiogenic characteristics. As studies have suggested a role for immune cells in immunosuppression in CMS4 (10–12), we examined the 1142 enriched gene sets in CMS4, and found 59 enriched gene sets that contained immune-related keywords.

We found an overlap of enriched immune-related gene sets between CMS1 and CMS4 (**Figure 1A**). These included production of Interleukin 6, cytokine secretion, and T-cell activation, all of which could lead to immune-induced cytotoxic activity that could destroy cancer cells, or chronic inflammation and escape in favour of cancer progression (3,36–38). Some gene sets had prominently higher enrichment scores in CMS1 compared to CMS4; among these was the gene set for T-cell activation. T-cell infiltration in CRC has been associated with better survival (39,40). We also found *Response to Molecule of Bacterial Origin* among these enriched gene sets, indicating the role microbes likely play in the characteristics of these subtypes.

**Figure 1.**
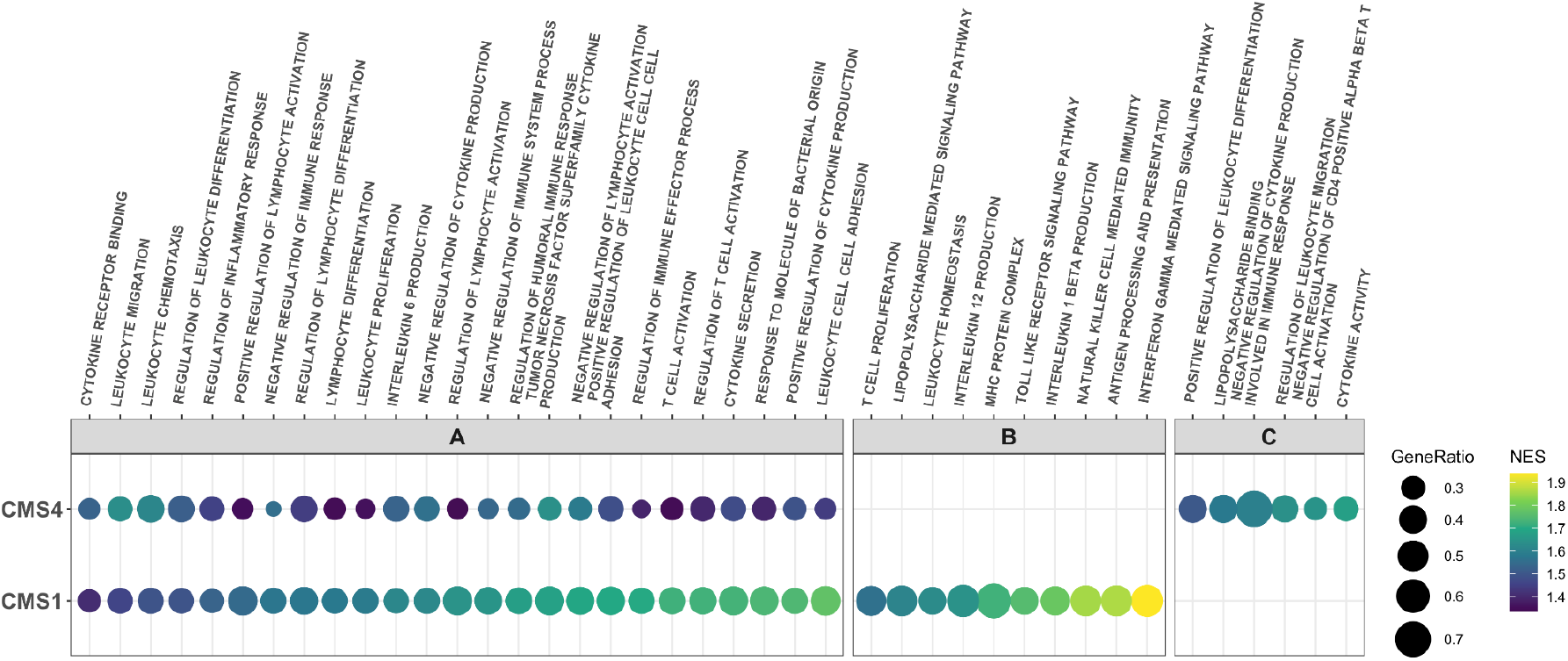
Immune Related Enriched Gene Sets in CMS1 and CMS4. **A.** Immune related enriched gene sets common to CMS1 and CMS4. **B.** Representative enriched immune-related gene sets unique to CMS1. **C.** Representative enriched immune-related gene sets unique to CMS4. *NES* = normalized enrichment score. *Gene Ratio* = genes contributing to enrichment of the gene set from our dataset divided by total set size of the gene set in question. All gene sets have adjusted *p-values < 0.05*.

We found 79 immune-related gene sets unique to CMS1. Several of these (e.g. antigen processing and presentation, MHC protein complex) indicate increased levels of antigen presentation, an early critical process in the induction of antitumor responses (41). Gene sets indicating activity of natural killer cells and production and response to interleukins, including Interleukin-12 and Interleukin-1 beta, were also unique to CMS1 (**Figure 1B**), in addition to a wide array of functions indicating regulation and homeostasis of immune responses, including regulation of apoptosis of leukocytes, lymphocytes, and T-cells. Genes associated with Toll-like receptor activity and lipopolysaccharide signaling pathways were also identified in CMS1, again suggesting that bacteria and their LPS may play a role in the responses observed.

CMS4 had 34 unique immune related enriched gene sets, several of which are associated with negative regulation of T-cells and other immune responses (**Figure 1C**). Lipopolysaccharide binding was also enriched in CMS4, suggesting a link to bacterial regulation, as seen in CMS1.

### Differentially abundant bacteria that contribute to LPS biosynthetic processes in CMS1 include Fusobacteria and Bacteroides fragilis species

From sequencing reads that did not map to the human genome, we obtained matches to microbial species. We found 296 microbial species that were differentially more abundant (DA) in CMS1 compared to the other CMS subtypes (p. adjust < 0.05). As our host GSEA had identified a ‘response to microbes’ among the enriched gene sets in CMS1 and CMS4, and that these may be related to LPS processes, we selected for bacteria that had proteins annotated with “Lipid A Biosynthetic Process” or “Lipopolysaccharide Biosynthetic Process” Gene Ontology terms, which identified 20 bacterial species in CMS1 (**Figure 2A, left**). Notably, we identified *Fusobacterium* and *Bacteroides* as among the abundant bacteria with LPS processes. These two genera have previously been implicated in the progression of CRC (17,22,23,42). We focused on *F. periodonticum* and *B. fragilis* to further investigate the potential interaction between their LPS molecules and immune responses in host cells (**Figure 2B**).

**Figure 2.**
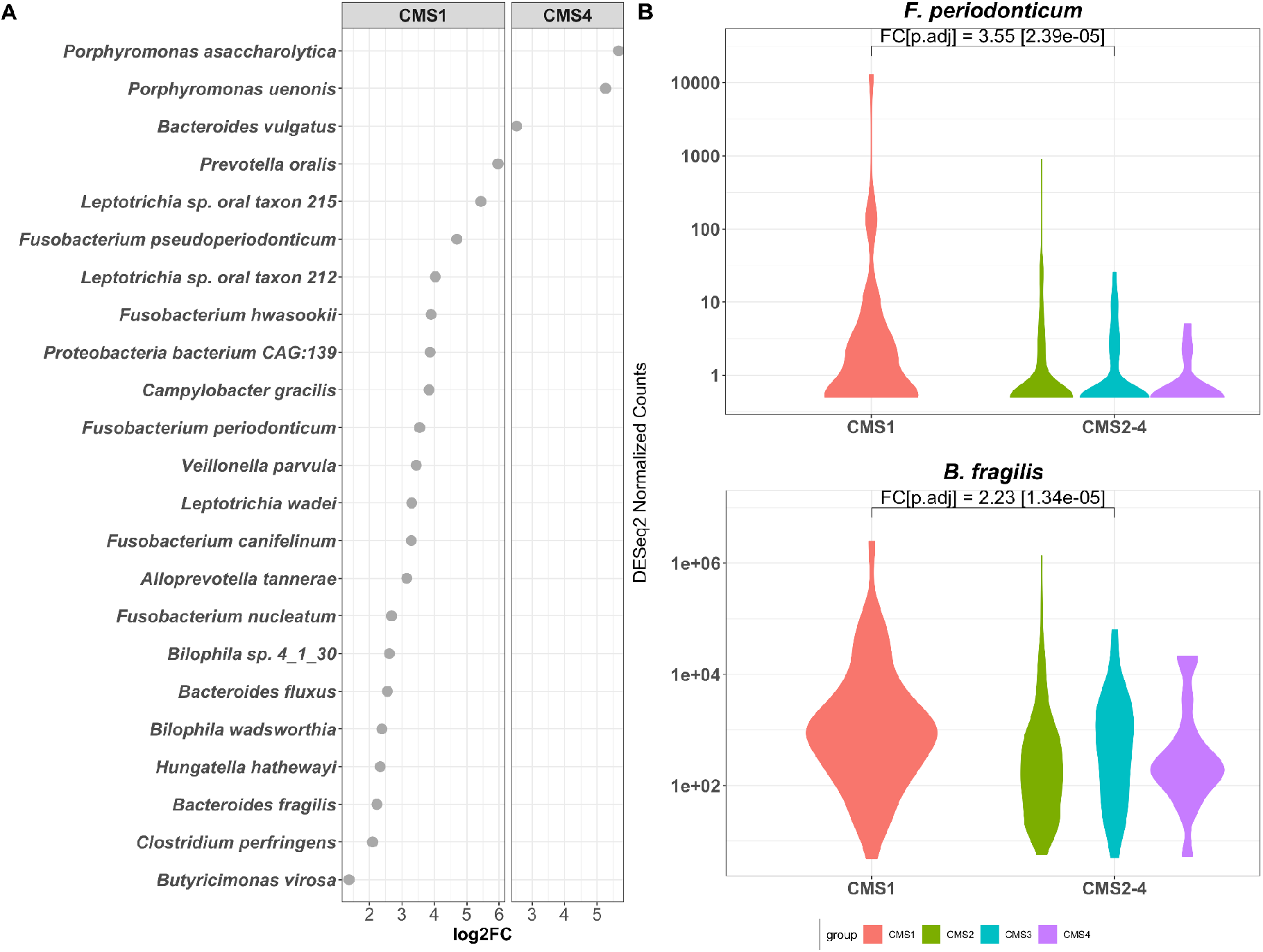
Abundant Microbes in CMS1 and CMS4 with lipopolysaccharide processes. **A.** Differentially abundant Bacteria in CMS1 and CMS4 with LPS processes annotations, showing their fold changes in CMS1 (left) or CMS4 (right) against the average abundance of the other three subtypes. **B.** *F. periodonticum* and *B. fragilis* normalized counts in CMS1 compared to the other subtypes. *log2FC =* log_2_Fold Change

In contrast, only 127 microbes were found to be differentially more abundant in CMS4 compared to the other three subtypes, and only three bacterial species had proteins annotated with LPS or Lipid-A biosynthetic processes (**Figure 2A, right**). Of these, we chose *Porphyromonas asaccharolytica* to investigate further for its effects on immune response in *in vitro* experiments, as *P. asaccharolytica* has been identified as a CRC marker in multi-cohort analyses (17,18).

### LPS from different bacterial species have different effects on cytokine release

Stock cultures of peripheral blood mononuclear cells (PBMCs) were incubated with increasing concentrations of LPS, extracted from *F. periodonticum (*strains *1/1/54 (D10), 2/1/31, and 1/1/41 FAA), B. fragilis (*strains *3/2/5, 2/1/16, and 2/1/56 FAA)*, or *P. asaccharolytica (*strains *CC44 001F*, and *CC1/6 F2)* overnight. The concentrations of cytokines present in the supernatants were then analysed by flow cytometry and compared to untreated PBMCs. We focused on IFN-γ, IL-6, IL-10, IL-12p70, IL-1β, and IL-18, as these were prominent cytokines identified in our gene-set enrichment analysis; IL-18 has been found to synergise with IL-12 to increase production of IFN-γ in T-cells (43) and IL-10 is known as a regulatory cytokine (37,38,44). All strains of a given species had similar effects on cytokine production.

The highest concentration of LPS tested from *B. fragilis* or *P. asaccharolytica* (600 ng/mL) was found to inhibit the release of the measured cytokines, compared to untreated PBMCs. Lower concentrations of LPS from these species also caused a decrease in cytokine production, but to a lesser degree (**Figures S1A and S1B**). Conversely, LPS from *F. periodonticum* strains showed stimulatory effects on the cytokines of interest, with low concentrations of LPS (6 ng/mL) causing an increase in their secretion compared to baseline PBMC levels (**Figure S1C**). No further increases were observed at higher LPS levels.

As all strains exhibited the same properties (**Figure S1**), we chose a single strain from each species, *F. periodonticum 2/1/31, B. fragilis 2/1/16*, and *P. asaccharolytica CC1/6 F2*, to use in subsequent experiments.

### B. fragilis and P. asaccharolytica LPS exhibit immunoinhibitory properties when co-cultured with stimulatory LPS from F. periodonticum

To further investigate the immune modulatory effects of different LPS, we tested whether LPS from *B. fragilis 2/1/16* or *P. asaccharolytica CC1/6 F2* could retain their immunoinhibitory effects on PBMCs when co-cultured with LPS from immunostimulatory *F. periodonticum 2/1/31*.

PBMCs were incubated with either *F. periodonticum* LPS alone, or in combination with either *B. fragilis* or *P. asaccharolytica* LPS, and cytokine production was measured. **Figure 3** and **Figure S2** show that the increased cytokine production observed in the presence of *F. periodonticum* LPS alone was attenuated by co-incubation with LPS from *B. fragilis* or *P. asaccharolytica*. In **Figure 3**, we show that *B. fragilis* or *P. asaccharolytica* LPS, when co-cultured with LPS from *F. periodonticum*, reduced cytokine secretion towards baseline levels in IL-18, IL-10, and IL-1β. These cytokines are considered protective against CRC, a regulatory cytokine, and detrimental in CRC, respectively. This reduction was significant for IL-1β, and IL-18 when *B. fragilis* LPS was co-incubated with *F. periodonticum* LPS (**Figure 3A and C**),and significant for all three cytokines when *P. asaccharolytica* LPS was co-incubated with *F. periodonticum* LPS (**Figure 3A, B, and C**). All significant comparisons had absolute effect sizes (Cohen’s d) greater than 0.8, indicating large effects (**Tables S1 and S2**).

**Figure 3.**
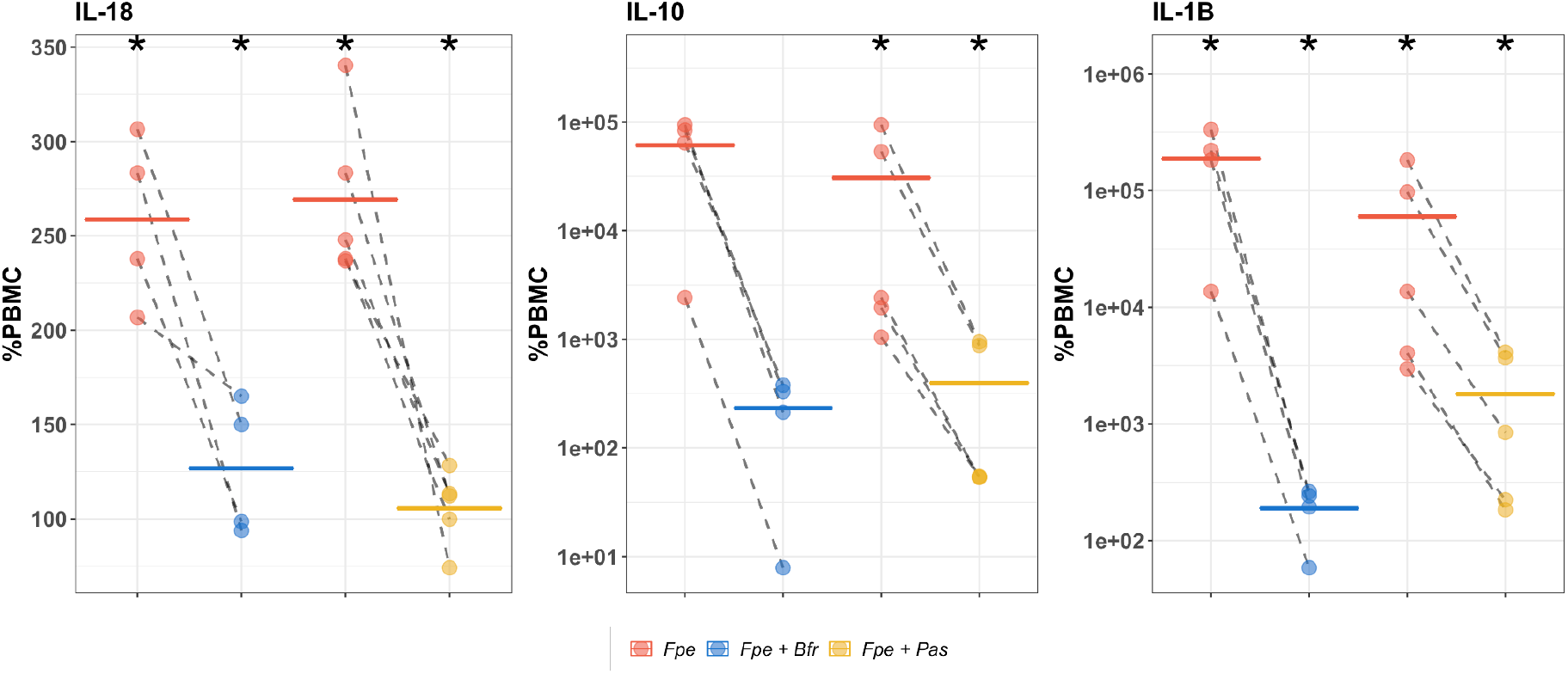
Changes in cytokine expression in peripheral blood mononuclear cells (PBMC) following treatment with *F. periodonticum* alone (red), or in combination with *B. fragilis* (blue) or *P. asaccharolytica* (yellow) for cytokines IL-18, IL-10, and IL-1β. For these experiments we used the lowest concentration of *F. periodonticum* LPS (6 ng/mL) and the highest concentration of LPS from *B. fragilis* or *P. asaccharolytica* (600 ng/mL), as these respective concentrations had the largest effects on cytokine production in earlier experiments. Values are shown as percentages of PBMC baseline secretion, which is set at 100%. Dashed lines indicate a single experimental run. Colored, solid horizontal lines represent the means of repeat experiments. Y-axes of IL-10 and IL-1β are in log_10_ scale, while Y-axis of IL-18 is in the linear scale. *Fpe = F. periodonticum* (6ng/mL), *Fpe + Bfr = F. periodonticum* (6ng/mL) +*B. fragilis* (600ng/mL), *Fpe + Pas = F. periodonticum* (6 ng/mL) +*P. asaccharolytica* (600 ng/mL)*; * = p-value < 0.05 (F. periodonticum vs PBMC, F. periodonticum + B. fragilis vs F. periodonticum* alone*, or F. periodonticum + P. asaccharolytica vs F. periodonticum* alone).

## Discussion

We combined gene-expression data from both tumor and associated microbiota to better understand how the microbiome could influence the differences seen in immune responses in CRC, in a way that could be tested *in vitro*. Several previous studies attributing function to the microbiome have largely been associative (17,18,45), and while important analytical results regarding functional contributions of the microbiome had been made, these have seldom been linked to host gene expression and functional contributions. Here, we show a potential link between enriched microbial species and immune responses seen in CRC, through the action of bacterial lipopolysaccharide.

Studies of immune responses in CRC have reported conflicting findings, where they can either induce tumor regression or lead to cancer progression. This delicate balance between pro- and anti-tumor effects is reflected in the consensus molecular subtypes (CMS) of CRC (5), with good prognosis CMS1 and poor prognosis CMS4 having immune infiltrates of differing compositions (10–12). Our analysis of host gene sets confirms previously published reports that CMS1 and CMS4 are immune infiltrated and inflamed, respectively, while CMS2 and CMS3 have little immune activation (35). Enriched gene sets unique to CMS1 involve antigen presentation and processing, natural killer cell activity, and homeostasis of immune responses, and regulation of T-cell apoptosis. These processes, as well as their regulation may affect how CMS1 tumors progress. Enriched gene sets in CMS4 meanwhile indicate an association with immunosuppression, consistent with previous studies (2,10)

Among the enriched immune gene sets common to CMS1 and CMS4, we identified those related to response to microbes and lipopolysaccharide, indicating a potential role for the microbiome in the characteristics of these subtypes. Focusing on bacteria in the tumor microenvironment with proteins annotated with LPS biosynthetic processes identified *F. periodonticum* and *B. fragilis* as differentially abundant in CMS1, and *P. asaccharolytica* differentially abundant in CMS4. *B. fragilis* was among the most enriched bacterial species in a multi-cohort analysis of CRC (17), while toxigenic strains of the bacteria are associated with immune activation, reactive oxygen species (ROS) production and DNA damage, as well as E-cadherin cleavage leading to cell proliferation (23,42). *Fusobacteria* species have been shown to associate with CRC in a multitude of studies (46) and *F. periodonticum* was reported to be enriched in MMR deficient (dMMR) tumors compared to MMR proficient (pMMR) tumors (16). *P. asaccharolytica*, meanwhile, has been identified as a CRC biomarker in previous studies (Dai et al 2018, Thomas et al 2019).

As CMS1 and CMS4 are thought to represent tumor-immune inhibitory and stimulatory environments, respectively, we hypothesized that *F. periodonticum* and *B. fragilis* would stimulate anti-tumor cytokines (IFN-γ, IL-12 IL-18), and decrease tumor-promoting cytokines (IL-1β and IL-6) (37,38), while the opposite effect may be seen with *P. asaccharolytica*. However, our results using human PBMCs indicate that *F. periodonticum* LPS stimulated the production of all cytokines of interest, including IL-10, while both *P. asaccharolytica* and *B. fragilis* inhibited the production of all of these cytokines. In addition, LPS from both *P. asaccharolytica* and *B. fragilis* could attenuate the immunogenicity of LPS from *F. periodonticum*, indicating that interactions between these microbes may add to the complexity of immune responses.

Although general attributes such as pro- or anti-tumor have been ascribed to many cytokines in the context of cancer, many cytokines display pleiotropic functions that may have opposing effects in CRC. For instance, decreased levels of IL-1β, along with IL-18, are correlated with increased colitis-associated cancer (CAC) in an inflammasome context (47,48) despite IL-18 being commonly viewed as anti-tumorigenic (37,38). IL-6 has anti-tumorigenic properties in the form of priming effector T-cells (36); and unchecked IFN-γ could compromise the colonic epithelial barrier (37,49) allowing an influx of microbiota that could influence cancer progression despite IL-6 being described as pro-tumorigenic, and IFN-γ being protective in CRC (37,38).

Our findings further illustrate that the balance and control in cytokine production is critical to determining whether the immune microenvironment is pro- or anti-tumorigenic. The balance, we hypothesise, may be reflected in the interactions we see in response to the bacterial LPS we have tested.

Although our results with *B. fragilis* LPS were unexpected, as high abundance of *B. fragilis* was associated with the immunogenic CMS1 tumors, previous studies have shown that LPS activity, conserved among the *Bacteroidales* order, can be immunosuppressive and promote immune tolerance to the high microbial load found in the gut, which hosts a complex microbial ecosystem (50). Indeed, species of the *Bacteroides* genus, including *B. fragilis*, have been found to have immunoregulatory properties (51–53).

Further, it was identified that enterotoxigenic *B. fragilis* could promote colonic tumors and induce inflammatory processes but not its non-toxigenic counterpart (23), indicating that it may be the toxin that is necessary for its inflammatory role and not LPS. We were unable to identify whether the *B. fragilis* in our CRC samples were toxigenic or non-toxigenic, or whether the toxigenic strains were expressing the BFT toxin; the BFT toxin may also only be produced within a certain timeframe during carcinogenesis (54). There is also a possibility of a mixture of non-toxigenic and toxigenic *B. fragilis* in our tumor samples, and this combination may indicate a nuanced balance between LPS and *B. fragilis* toxin, along with the more immunogenic LPS of other microbes such as *F. periodonticum*.

Our findings indicate that while LPS from *B. fragilis* may be immunosuppressive, interaction with immunogenic molecules produced by the same or other microbes adds a layer of complexity to the balance of immune responses in colorectal cancer. In CMS1, we postulate that this balance skews towards immune activation, contributing to anti-tumor effects and positively affecting prognosis, while the immunosuppressive LPS of *P. asaccharolytica* may contribute to immune evasion and escape of tumors in CMS4.

The cytokine release-inhibiting capabilities of LPS from *B. fragilis* and *P. asaccharolytica* are notable as previous studies emphasize the pro-inflammatory activities of CRC-associated microbiota (13,23,55–57). While we acknowledge that these events also occur within our tumor samples, we also suggest that microorganisms play a role in immunosuppression, either by aiding the tumor progression through immune evasion and escape, or contributing to homeostasis of immunogenic processes.

The limitations of the study include the use of PBMCs as a proxy for immune cells in the tumor microenvironment. PBMCs may not adequately reflect the effects of LPS on tumor-infiltrating lymphocytes. In vitro cultures of single cell types do not allow for crosstalk between different cell populations, which would be expected in the complex tumor microenvironment. Furthermore, we acknowledge that while we have tested the effects of LPS from single species and pairs of species, colorectal tumors may harbour up to hundreds of different species, and each may elicit an effect dependent on LPS structure and absolute bacterial counts, which could contribute to the nuanced immune-modulation within the TME. In addition, the effects of LPS may be countered or exacerbated by other known bacterial mechanisms, e.g. bacterial toxins, or as yet undiscovered interactions.

## Conclusion

In this study, we identified gene sets involved in immune response to microbial triggers in both CMS1 and CMS4 colorectal cancer subtypes and identified LPS from particular bacterial species that associate with immune responses. In vitro analyses showed that *F. periodonticum* LPS, found in CMS1 tumors, increased production of cytokines IL-1β, IFN-γ, IL-18, IL-10, IL-6, and IL-12p70, while LPS from *B. fragilis*, found in CMS1 tumors, and *P. asaccharolytica* LPS, found mainly in CMS4 tumors, decreased production of these cytokines and could also attenuate the immunogenic effect of *F. periodonticum* LPS. Where most previous studies focus on the inflammation-inducing capabilities of CRC-associated microorganisms, our results indicate that their immunosuppressive potential should not be overlooked and adds another layer of complexity to immune responses in CRC.

## Declarations

## Acknowledgements

The authors would like to thank Helen Morrin at the Cancer Society Tissue Bank, Christchurch, and the patients involved and their whānau for generously participating in this study.

## Funding

Maurice and Phyllis Paykel Trust

Gut Cancer Foundation (NZ), with support from the Hugh Green Foundation

Colorectal Surgical Society of Australia and New Zealand (CSSANZ)

The Health Research Council of New Zealand

The funding bodies had no role in the design of the study or collection, analysis, and interpretation of data or in writing the manuscript.

