## Supplementary Material for "Bacterial lipopolysaccharide modulates immune response in the colorectal tumor microenvironment"

**SUPPLEMENTARY TABLES**

| **Table S1. Test of Differences between Cytokine Concentrations Released after PBMC Treatment using *F. periodonticum and B. fragilis* LPS** | | | | | | | |
|---|---|---|---|---|---|---|---|
| **Cytokine** | **Reference Group** | **Group 2** | **N1** | **N2** | **p** | **Cohen's d** | **Magnitude** |
| **IL-12p70** | PBMC | Fp6 | 3 | 3 | 0.16 | -0.72 | moderate |
| | Fp6 | Fp6 + Bf600 | 3 | 3 | 0.20 | 0.62 | moderate |
| **IL-10** | PBMC | Fp6 | 4 | 4 | 0.057 | -1.10 | large |
| | Fp6 | Fp6+Bf600 | 4 | 4 | 0.055 | 1.11 | large |
| **IFN-γ** | PBMC | Fp6 | 4 | 4 | 0.09 | -0.90 | large |
| | Fp6 | Fp6+Bf600 | 4 | 4 | 0.09 | 0.90 | large |
| **IL-1β** | PBMC | Fp6 | 4 | 4 | 0.041 | -1.26 | large |
| | Fp6 | Fp6+Bf600 | 4 | 4 | 0.041 | 1.26 | large |
| **IL-18** | PBMC | Fp6 | 4 | 4 | 0.017 | -1.77 | large |
| | Fp6 | Fp6+Bf600 | 4 | 4 | 0.05 | 1.163 | large |
| **IL-6** | PBMC | Fp6 | 4 | 4 | 0.025 | -1.52 | large |
| | Fp6 | Fp6+Bf600 | 4 | 4 | 0.023 | 1.57 | large |
| **Legend:** Fp6=*F. periodonticum 2/1/31* (6 ng/mL) alone; Fp6+Bf600 =*F. periodonticum 2/1/31* (6 ng/mL) + *B. fragilis 2/1/16* (600 ng/mL) co-incubation; N=number of repeats used in statistical calculations; p=paired t-test p-value; d=cohen's d; Magnitude = interpretation of cohen's d | | | | | | | |

| **Table S2. Test of Differences between Cytokine Concentrations Released after PBMC Treatment using *F. periodonticum* and *P. asaccharolytica* LPS** | | | | | | | |
|---|---|---|---|---|---|---|---|
| **Cytokine** | **Reference Group** | **Group 2** | **N1** | **N2** | **p** | **Cohen's d** | **Magnitude** |
| **IL-12p70** | PBMC | Fp6 | 4 | 4 | 0.06 | -1.07 | large |
| | Fp6 | Fp6+Pa600 | 4 | 4 | 0.08 | 0.95 | large |
| **IL-10** | PBMC | Fp6 | 5 | 5 | 0.017 | -1.41 | large |
| | Fp6 | Fp6+Pa600 | 5 | 5 | 0.016 | 1.45 | large |
| **IFN- γ** | PBMC | Fp6 | 4 | 4 | 0.035 | -1.34 | large |
| | Fp6 | Fp6+Pa600 | 4 | 4 | 0.06 | 1.07 | large |
| **IL-1β** | PBMC | Fp6 | 5 | 5 | 0.010 | -1.63 | large |
| | Fp6 | Fp6+Pa600 | 5 | 5 | 0.011 | 1.61 | large |
| **IL-18** | PBMC | Fp6 | 5 | 5 | 0.02 | -1.35 | large |
| | Fp6 | Fp6+Pa600 | 5 | 5 | 0.021 | 1.33 | large |
| **IL-6** | PBMC | Fp6 | 3 | 3 | 0.017 | -2.52 | large |
| | Fp6 | Fp6+Pa600 | 3 | 3 | 0.015 | 2.69 | large |
| **Legend:** Fp6=*F. periodonticum 2/1/31* (6 ng/mL) alone; Fp6+Pa600=*F. periodonticum 2/1/31* (6 ng/mL) + *P. asaccharolytica CC1/6 F2*(600 ng/mL) co-incubation; N=number of repeats used in statistical calculations; p=paired t-test p-value; d=cohen's d; Magnitude = interpretation of cohen's d | | | | | | | |

### SUPPLEMENTARY FIGURES

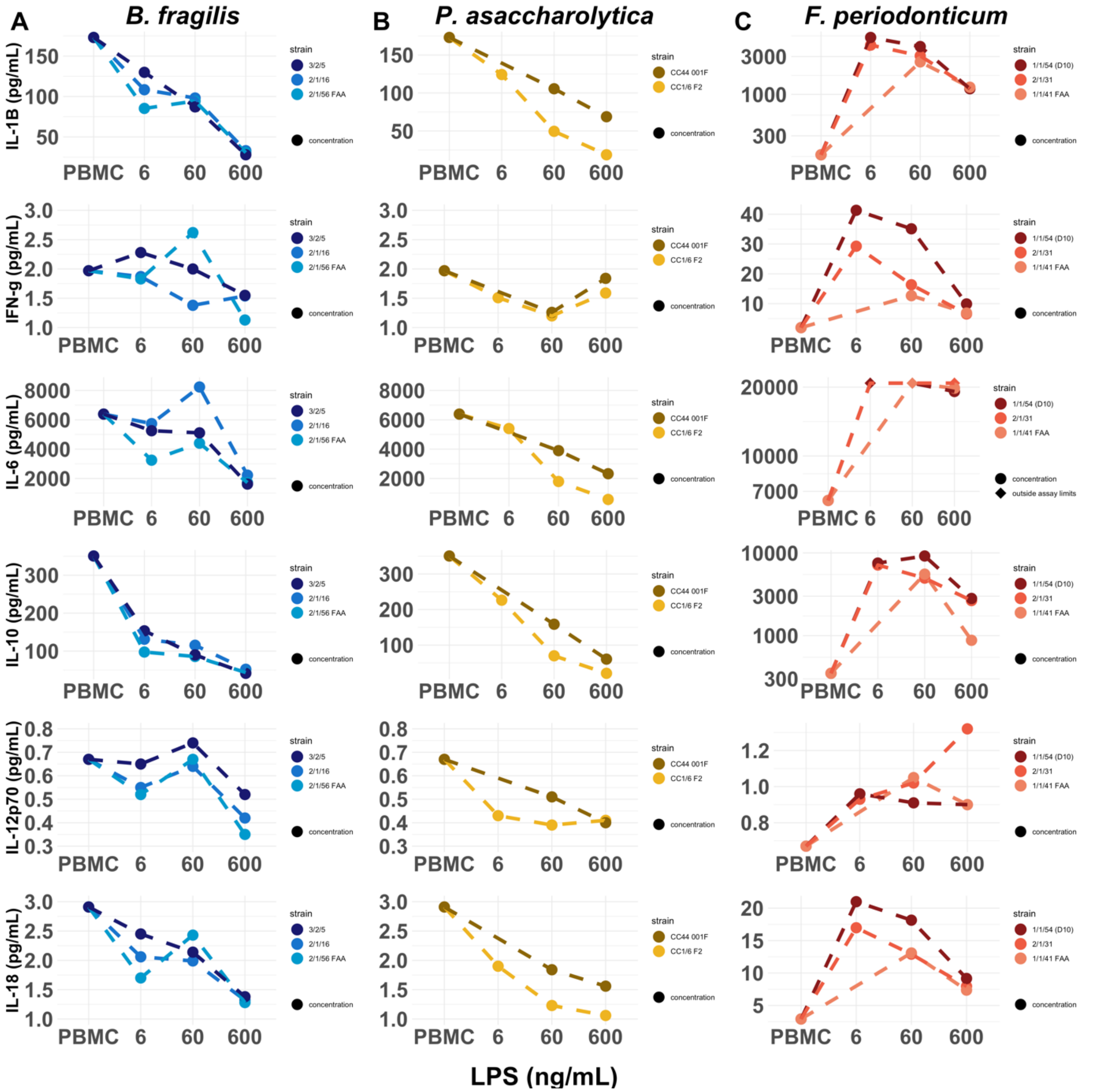

**Figure S1. Secreted cytokine concentrations (pg/mL) after overnight incubation of PBMCs with different LPS concentrations (6 ng/mL, 60 ng/mL, 600 ng/mL) from different strains of A. *B. fragilis* B. *F. periodonticum*, and C. *P. asaccharolytica* compared to PBMC baseline (no treatment)**. Dashed lines connect data points from the same strains. Colors differentiate species (*B. fragilis:* blue; *P. asaccharolytica*: yellow; *F. periodonticum:* red) and different shades of a color differentiate the strains of each species. Where values are outside of assay limits, we show the assay limit, and denote the point with '◆'. For *F. periodonticum* plots, IL-1β, IL-6, and IL-10 y-axes are at scale $\log_{10}$.

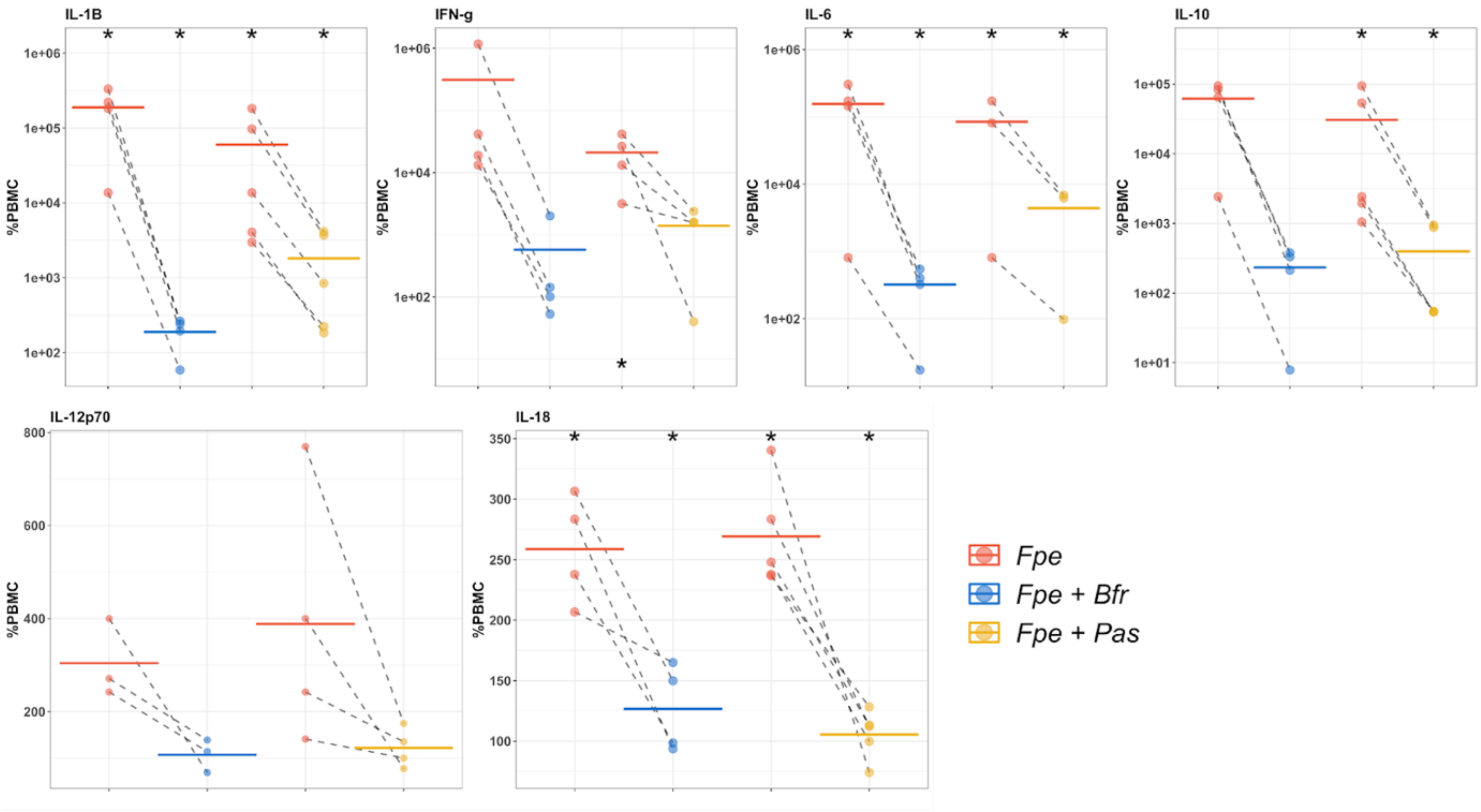

**Figure S2. Changes in cytokine expression in peripheral blood mononuclear cells (PBMCs) following treatment with *F. periodonticum* alone (red) or in combination with *B. fragilis* (blue) or *P. asaccharolytica* (yellow).** We show results for cytokines of interest IL-1β, IFN-γ, IL-6, IL-10, IL-12p70, and IL-18. *F. periodonticum 2/1/31* LPS used was at a concentration of 6 ng/mL**.** *B. fragilis 2/1/16* LPS and *P. asaccharolytica CC1/6 F2* LPS used were at a concentration of 600ng/mL. Values are shown as percentages of PBMC baseline secretion, which is set at 100%. Dashed lines indicate a single experimental run. Colored, solid horizontal lines represent the means of repeat experiments. Y-axes of IL-1β, IFN-γ, IL-6, and IL-10 are in $\log_{10}$ scale, while Y-axes of IL-12p70 and IL-18 are in linear scale.

*Fpe = F. periodonticum* (6ng/mL), *Fpe + Bfr = F. periodonticum* (6ng/mL) + *B. fragilis* (600ng/mL), *Fpe + Pas = F. periodonticum* (6 ng/mL) + *P. asaccharolytica* (600 ng/mL)*; (*) = p-value < 0.05 (F. periodonticum vs PBMC, F. periodonticum + B. fragilis vs F. periodonticum* alone*, or F. periodonticum + P. asaccharolytica vs F. periodonticum* alone*).*
